# Understanding short-term interactions between arousal and attention in infants and children: applying the Aston-Jones framework

**DOI:** 10.1101/190108

**Authors:** Sam V. Wass

## Abstract

Differential Susceptibility Theory explains long-term associations between neurobiological sensitivity and cognitive outcomes, but no comparable theoretical framework exists to understand how neurobiological sensitivity and cognitive performance inter-relate on shorter time-frames. Here, we evaluate a framework proposed by Aston-Jones and colleagues, building on the Yerkes-Dodson model, to understand these associations. The framework describes how slow-moving (tonic) changes in autonomic arousal relate to fast (phasic) changes, as observed for example relative to experimenter-determined events, and how phasic changes relate to attention. Larger phasic changes, which associate with better selective attention, are most likely at mid-level tonic arousal. Smaller phasic changes, and worse selective attention, are observed at hypo-and hyper-arousal. We review the fit of this model to typical and atypical development, during infancy and childhood.

## Introduction

### Measuring autonomic nervous system function in infants and children

The Autonomic Nervous System (ANS) serves as the fast-acting, neural substrate of the body’s stress response (e.g. Cacioppo, Tassinary, & Berntson, 2000; Tsigos & Chrousos, 2002; Ulrich-Lai & Herman, 2009). It acts in concert with the Hypothalamic-Pituitary-Adrenal (HPA) axis, which is the slow-acting, endocrine substrate of stress (Cacioppo et al., 2000). Relationships between the ANS and HPA axis are complex (Kolacz, Holochwost, Gariepy, & Mills-Koonce, 2016; Quas et al., 2014) and a full discussion of them is beyond the scope of the present paper. In this article, arousal is defined as referring to the total levels of activity within the ANS.

The ANS is thought to operate primarily via the norepinephrine and cholinergic neurotransmitters, and to be governed by a range of brain areas centered on the locus coeruleus (LC) and reticular pathways in the brainstem, that communicate via thalamic and cortical projections (e.g. Amaral & Sinnamon, 1977; Arnsten & Goldman-Rakic, 1984). It acts through two complementary systems - the sympathetic (SNS) and parasympathetic (PNS) nervous systems (Cacioppo et al., 2000). The sympathetic nervous system (SNS) is involved in quick response mobilizing (‘fight or flight’) responses. The parasympathetic nervous system (PNS) is involved in more slow-acting and response dampening (‘rest or digest’) responses (Ulrich-Lai & Herman, 2009). In many cases the SNS and PNS have opposite effects, although their function is non-additive (Berntson et al., 1994; Lacey, 1967).

In animal research, ANS activity is usually indexed by recording from the LC in the brainstem (Aston-Jones & Cohen, 2005; Ulrich-Lai & Herman, 2009). In humans this is not possible, and so research into the ANS generally uses one of several indirect measures. Heart Rate (HR) is thought to receive contributions from both SNS and PNS, with faster HR indexing greater SNS and less PNS activity (McCabe, Schneiderman, & Field, 2000). Respiratory Sinus Arrhythmia (RSA), derived from HR, indexes the degree to which heart rate changes relative to respiratory cycles, and is thought to index PNS activity (Anrep, Pascual, & Rossler, 1935). Higher RSA (also known as increased heart rate variability, or increased vagal tone) indexes more PNS activity (Anrep et al., 1935). Impedance cardiography (ICG) measures the Pre-Ejectile Period, the time interval between the heart beat and the outflow of blood from the aorta, and is thought to index SNS activity (Oberlander, Grunau, Pitfield, Whitfield, & Saul, 1999). A longer Pre-Ejectile Period indexes more SNS activity (Oberlander et al., 1999). Electrodermal Activity (EDA, also known as Galvanic Skin Response) is thought to measure SNS control (Shields, Macdowell, Fairchild, & Campbell, 1987). Higher EDA indexes more SNS activity. Pupil size is thought to be influenced by both SNS and PNS. A larger pupil size indexes more SNS and less PNS (Loewenfeld, 1993). Body movement is often traditionally included in operational definitions of arousal (Pfaff, 2005). In addition, ERP components (such as the P3 fronto-parietal component) (Sutton, Braren, Zubin, & John, 1965), frequency spectral components (theta/beta ratios and alpha fluctuations) (see Imeraj et al., 2012) and short bursts of high frequency brain activity known as micro-arousals (Sasai, Matsuura, & Inoue, 2013; Sforza, Jouny, & Ibanez, 2000) have all been described as neural indices of arousal. However, for reasons of space, these neural measures have not been included the present paper.

Most of the research included in this review only reports data from one of these peripheral indices, leaving considerable challenges to researchers wishing to integrate findings across methods (Lacey, 1967; Loewenfeld, 1993; Sanders, 1983; Taylor & Epstein, 1967). To address this, recent research studied tonic and phasic covariation between HR, EDA, pupil size and body movement patterns in typical 12-month-old infants (Wass, Clackson, & de Barbaro, 2016; Wass, de Barbaro, & Clackson, 2015). It was found that HR and body movement patterns showed strong and reliable associations. Associations with EDA were consistently observed, but appeared to be specific to moments of high general arousal. Tonic pupil size was found to associate with tonic levels of HR and movement, but not with EDA.

Arousal levels are in a continuous state of slow-moving flux. Our primary periodic arousal change is a circadian one (Silver & LeSauter, 2008). Circadian rhythms determine firing rates of Norepinephrine neurons in the rat LC (e.g. Aston-Jones & Bloom, 1981a) and also determine changes in ANS activity and PNS/SNS balances (Baehr, Revelle, & Eastman, 2000). Cyclical fluctuations in arousal at smaller, sub-hour scales have also been observed in fetuses and infants (Feldman, 2006; Robertson, 1993). They may also exist on large time-scales, such as annual cycles – although this question has not, to our knowledge, been investigated.

A variety of other, non-cyclical factors have also been shown to influence arousal. In animals mild, experimentally controlled stressors such as an air puff (S. J. Grant, Aston-Jones, & Redmond, 1988) and other environmental (Abercrombie & Jacobs, 1987) or physiological stressors (Morilak, Fornal, & Jacobs, 1987) have been shown to lead to increases in LC activity. In humans, mild stressors such as an arm restraint procedure (Garcia-Coll et al., 1988) or a ‘still face’ manipulation (K. A. Grant et al., 2009) have all been shown to lead to increases in arousal in infants and children. Other types of experimenter-controlled events, such as new, attention-getting stimuli, have also been shown to lead to decreases in arousal (Richards, 2010; Sokolov, 1963). In addition, purely internal triggers such as thoughts (realizing that one has lost one’s keys) are also considered likely to lead to phasic changes in arousal, although these types of internal triggers have not been studied in detail (cf. Stawarczyk, Majerus, Maj, Van der Linden, & D’Argembeau, 2011).

### Long-term associations between arousal, arousal reactivity and cognitive outcomes

Differential Susceptibility Theory (DST) suggests that some individuals are more susceptible than others to *both* negative (risk-promoting) and positive (development-enhancing) environmental conditions (Belsky, Bakermans-Kranenburg, & van IJzendoorn, 2007; Boyce & Ellis, 2005; Ellis, Essex, & Boyce, 2005). Individuals with high biological sensitivity to context show superior long-term outcomes in positive environments, but worse long-term outcomes in negative environments; children with lower sensitivity show a lesser influence of environment on long-term outcomes.

Researchers working within the theoretical framework of this model have used a variety of techniques to index biological sensitivity. First, some have examined the *genetic* factors that influence an individual’s level of sensitivity to their environment (e.g. Belsky et al., 2007; Belsky et al., 2009; Bakermans-Kranenburg & van Ijzendoorn, 2011). For example, children with less efficient dopamine-related genes were found to perform worse in negative environments than comparison children without the ‘genetic risk’, but they also profited most from positive environments (Bakermans-Kranenburg & van Ijzendoorn, 2011). In genetics, biological sensitivity is typically conceptualized in terms of plasticity (Belsky et al., 2009). Relationships are investigated over long-term time-scales, in order to examine the life-long relationship between genetics, environment and cognitive and behavioral outcomes. However, the question of how these ‘genetically sensitive’ individuals differ over shorter time-frames, such as on a trial-by-trial basis, remains relatively under-explored.

Second, another body of research has used questionnaire-based assessments of *temperament* (Rothbart, 2007) to index an individual’s level of biological sensitivity to context. For example, a meta-analysis found that children with high negative affect were found to be more sensitive to parenting style than others, although associations were only present when the trait was assessed during infancy (Slagt, Dubas, Deković, & van Aken, 2016). Again, relationships tend to be investigated over life-long time-scales, but how these individuals differ in performance over shorter time-frames is a question that has received relatively little research (although see e.g. Sheese, Rothbart, Posner, White, & Fraundorf, 2008).

A third body of research has used *physiological reactivity* to index biological sensitivity to context (see Obradovic, 2016 for a review). Physiological reactivity is usually assessed by measuring the degree of ANS (HR or RSA) or HPA axis (cortisol) change to an experimental stressor in the lab (see Burt & Obradovic, 2013). Environment is assessed, for example, via life histories, questionnaires, or video-coding of parent-child interactions. A variety of types of long-term outcome have been investigated, such as physical and mental health (e.g. Boyce et al., 1995), and socioemotional and cognitive competencies, but an increasing number of studies are focusing on outcomes such as academic performance or performance on executive function tasks (Obradovic, 2016). Thus, for example, Skowron and colleagues examined RSA responses to a social challenge task in 161 children aged 3-5 years, of whom half had been exposed to childhood maltreatment. To the same children they also administered experimental assessments of inhibitory control, using age-appropriate Stroop tasks. They found that elevated parasympathetic reactivity during a parent–child interaction was associated with less optimal performance on EF tasks in 3- to 5-year-olds exposed to maltreatment, and with more optimal performance among non-maltreated children (Skowron, Cipriano-Essel, Gatzke-Kopp, Teti, & Ammerman, 2014; see also e.g. Obradovic, Bush, Stamperdahl, Adler, & Boyce, 2010; Spinrad et al., 2009; Ursache, Blair, Stifter, Voegtline, & Investigators, 2013).

Again, the majority of research has investigated *life-long* consequences of an individual’s level of physiological reactivity. The question of how more sensitive, or reactive, individuals show performance differences over shorter time-frames, such as on a trial-by-trial basis, has received relatively little attention.

The ANS, however, shows change on sufficiently fine-grained time-scales that change can be measured on a trial-by-trial basis. A number of studies using convergent methodologies have shown that individual trials that show greater trial-by-trial ANS responses show better performance (see e.g. Schoen, Miller, Brett-Green, & Hepburn, 2008; Richards, 1997b). This suggests a question: do the same individuals who show a large arousal change to a negative stimulus (mild experimental stressors such as arm restraint) also show a large arousal change to a positive stimulus (such as a new item to be memorized in a working memory task, or a novel image in a habituation task)? If so, this might be a mechanism by which biological sensitivity to context might confer an advantage in certain situations, and a disadvantage in others. Addressing this question is one of the main aims of the present paper.

Another point that currently appears under-specified is the question of whether physiological reactivity is a static, time invariant, aspect of individual differences, or whether reactivity changes over time. Most research in this area tends to assume the former; however, the latter possibility appears intuitive: if I am tense or worried and someone behind me drops a plate, my reaction will be greater than at other times. No theoretical framework exists, however, for incorporating this kind of short-term variability within the context of DST.

### The Aston-Jones model (AJM)

Yerkes and Dodson first suggested that the relationship between arousal and performance is curvilinear, such that the highest levels of performance are observed at intermediate levels pf physiological or mental arousal (Yerkes & Dodson, 1908). More recent research from Aston-Jones and colleagues has developed this model (Usher, Cohen, Servan-Schreiber, Rajkowski, & Aston-Jones, 1999).

Aston-Jones and colleagues distinguished between two modes of LC function. In a phasic mode, reactive bursts of LC activity are associated with the outcome of task-related decision processes and are closely coupled with behavioral responses. In a tonic mode, levels of baseline LC activity follow complex patterns of spontaneous change in response to cyclical and environmental factors that are not closely coupled with behavioral responses (Aston-Jones & Bloom, 1981b; S. J. Grant et al., 1988).

The phasic and tonic properties of LC firing are thought to be inversely related to one another by means of a single parameter: electrotonic coupling among LC neurons (Usher et al., 1999; Aston-Jones & Cohen, 2005). Extreme high and low levels of tonic activity were found to associate with fewer phasic LC responses, but mid-level tonic activity was associated with large phasic responses (see Figure 1). This is consistent with the ‘inverted U’ relationship predicted by Yerkes and Dodson (Yerkes & Dodson, 1908). The same research also found that phasic LC responses are more tightly linked temporally to the behavioral response than to the sensory stimulus, suggesting that phasic LC responses are driven by decision processes rather than by sensory or motor activities *per se* (Usher et al., 1999).

**Figure 1:**
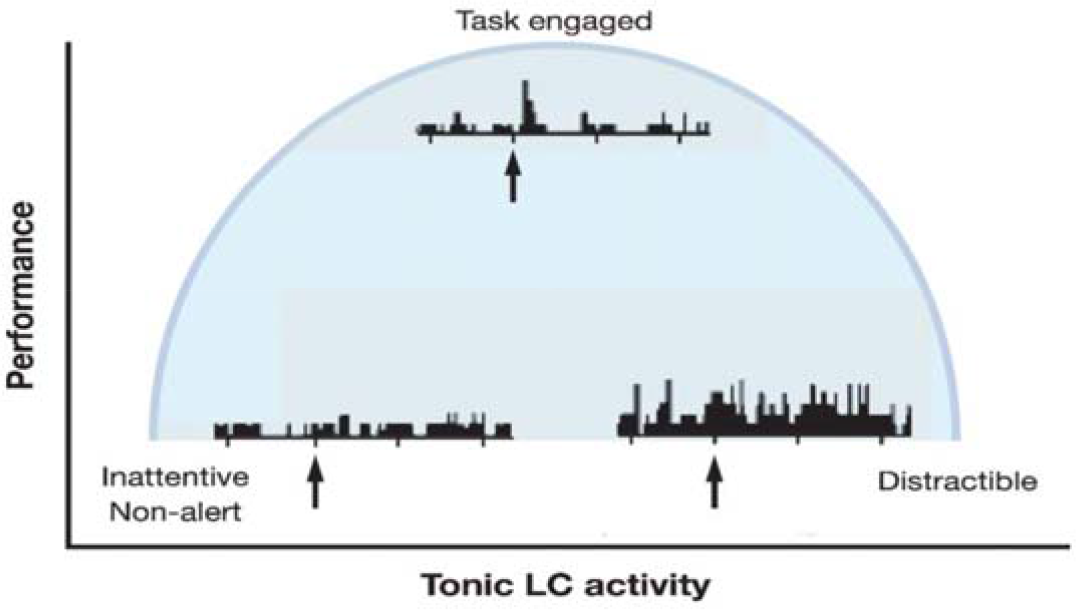
Schematic from Aston-Jones and Cohen. At mid-level Tonic LC activity, phasic (event-related) LC changes are greater and the capacity to maintain focused attention is increased. At both hypo-and hyper-tonic arousal, phasic LC changes are reduced and the capacity to maintain focused attention is lower (Figure from Aston-Jones & Cohen, 2005).

According to the Aston-Jones model (AJM), high tonic LC firing rates associate with low phasic responsiveness (Aston-Jones & Cohen, 2005). Individuals with high tonic LC are more vigilant or stimulus driven, and show better signal-to-noise ratios in primary cortices, increased habitual responding, increased fear conditioning and increased memory consolidation (Aston-Jones, Rajkowski, & Kubiak, 1997; Rajkowski, Majczynski, Clayton, & Aston-Jones, 2004). Mid-level tonic firing is associated with increased phasic responsiveness, together with superior working memory and selective attention with task irrelevant cues. Low-level tonic firing is associated with general inattentiveness, and reduced phasic responsiveness. Of note, the model is agnostic as to whether the phasic changes observed to stimulus events should be short-term *increases*, or *decreases*, in ANS – predicting merely that a change should take place and that a larger stimulus-related change associates with better performance.

The AJM is consistent with research from other disciplines into the effect of stress on attentional behaviors in humans. Neuroimaging suggests that high levels of stress are associated with down-regulation of areas such as the dorsolateral pre-frontal cortex that are involved in directed attention, together with up-regulation of areas including the hypothalamus, striatum, amygdala and occipital cortices involved in bottom-up, salience-driving orienting (Arnsten, 2009). At lower levels of stress (analogous to Aston-Jones’ mid-level tonic arousal), frontal cortical areas show increased activity (Arnsten, 2009). Behaviorally this leads to a phenotype in which high stress is associated with decreased voluntary control of attention and increased oculomotor responsivity to salient peripheral targets, whereas lower stress is associated with increased voluntary control (Alexander, Hillier, Smith, Tivarus, & Beversdorf, 2007; Broadbent, 1971). However, the current models from cognitive neuroscience do not explore the neural correlates of hypo-arousal, as predicted by the Yerkes-Dodson and Aston-Jones models. Presumably these would include globally decreased task-related activation.

Murphy and colleagues examined the predictions of the Aston-Jones model in adult participants (Murphy, Robertson, Balsters, & O’Connell, 2011). They tested performance on an auditory oddball task that required participants to detect 500Hz tones interspersed at a rate of 20% among 1000Hz tones. To index tonic arousal, they measured spontaneous changes in pupil size. They also recorded phasic, attention-related changes in the P3 ERP component locked to each stimulus presentation. The P3 has been associated with LC-mediated ANS changes (Nieuwenhuis, De Geus, & Aston-Jones, 2011). They found that both P3 amplitudes and task performance (reaction times) showed an inverted U-shaped relationship, such that largest P3 amplitudes and optimal performance occurred at the same intermediate level of pupil diameter (Murphy et al., 2011).

Of note, the AJM defines high and low levels of tonic activity only relative to the average activity level *for that individual*. The model does not explicitly deal with between-participant variance: the question of whether individuals who have higher, or lower, average arousal levels *relative to other individuals* tend to show greater phasic responsiveness, or different attentional profiles. For the purposes of this article we have assumed that the model makes equivalent predictions for understanding between-as within-participant variance, as we describe below.

Therefore, the model makes the following predictions:

i. hypo-aroused individuals (who show low average arousal levels relative to others) should show worse voluntary attention control (decreased general attentiveness) and reduced phasic arousal changes to stimulus events.
ii. ii) mid-level aroused individuals should show superior voluntary attention control and increased phasic arousal changes to stimulus events.
iii. hyper-aroused individuals should show worse voluntary attention control (increased distractibility) and reduced phasic arousal changes to stimulus events.
iv. the same features should be observed when considering variance within individuals as between individuals – i.e. considering variance observed within a particular individual across the course of a day, or within a particular testing session.
v. larger phasic arousal changes to stimulus events should be associated with better stimulus encoding.

In the following sections we review the available evidence in infancy (part 2) and childhood (part 3), and evaluate their goodness of fit to this model. First (part 1) we address some preliminary questions concerning how tonic (baseline) and phasic arousal reactivity changes over the course of development.

### Part 1 - Preliminary questions

In this section, prior to addressing the specific predictions of the AJM, we first address three more basic questions. First, do ongoing fluctuations in arousal associate with fluctuations in children’s voluntary attention control capacities? Answering this question is vital to motivate the biological approach that we have taken to understanding differences in cognition. Second, how does tonic (or baseline) arousal change over the course of typical development? Third, how do phasic (or reactive) changes in arousal to stimulus events develop with increasing age? This is essential to provide ‘norming’ data, and in order to contextualize other types of changes, such as behavioral changes over time.

#### 1) Do ongoing fluctuations in arousal associate with fluctuations in cognitive performance?

The AJM predicts that spontaneous fluctuations in arousal should co-vary with fluctuations in individuals’ voluntary attention control. Convergent research has supported this prediction (Bacher & Robertson, 2001; see also Richards, 2010, Richards, 2010, 2011). de Barbaro and colleagues collected continuous data in 20-minute segments while presenting a battery of mixed novel static and dynamic viewing data to a cohort of typical 12-month-old infants (de Barbaro, Clackson, & Wass, in press). They measured attention by recording the duration of infants’ individual looks to the screen, and arousal by recording a composite of HR, EDA and movement (see Figure 2). They found that fluctuations in arousal related to fluctuations in attention, such that increased arousal was associated with shorter look durations. Using cross-correlations they showed that these associations disappeared when a time lag of more than 100 seconds was introduced between arousal and attention, suggesting that they are temporally specific. Finally, also using cross-correlations, they found that arousal levels up to 25 seconds before the onset of a look were predictive of the duration of that look, whereas the reverse was not true (de Barbaro et al., in press; Wass et al., 2016; see also Bacher & Robertson, 2001). This suggests that changes in arousal tend to precede subsequent changes in look duration.

**Figure 2:**
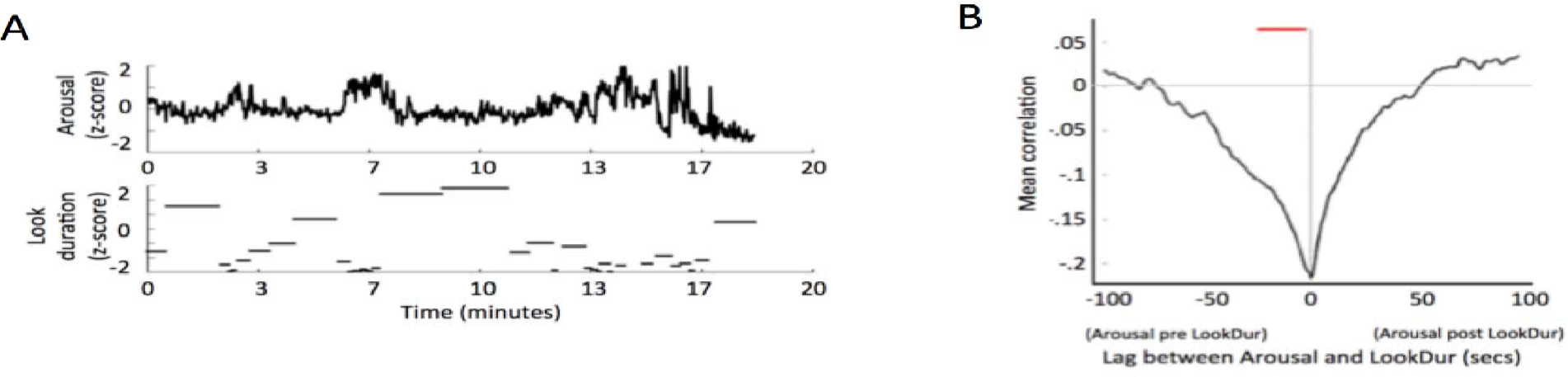
A) sample of raw data from the study from de Barbaro and colleagues. Infants were presented with a 20-minute battery of mixed static and dynamic viewing materials, and their changing arousal levels (top plot) and durations of looks to the stimulus presentation area (bottom plot) were measured. B) Cross-correlation plot showing how the relationship between arousal and look duration changes as a function of varying the time-interval between them. X-axis indicates the time-lag between the two variables. Y-axis indicates the correlation observed between the two variables, when one was time-lagged relative to the other. Colored dots above the plot indicate whether a significant relation was observed at each time-lag p<.05). Three aspects of the results can be seen. First, at time 0 (no lag), a negative relationship can be seen, suggesting that, at moments of high arousal, look durations tend to be lower. Second, this relationship disappears when a lag of more than 100 seconds is introduced between the two variables, suggesting that it is temporally specific. Third, the lag is asymmetric: arousal levels up to 25 seconds before the onset of a look were predictive of the duration of a look, whereas the reverse was not true. This suggests that changes in arousal tend to precede changes in look duration. Figures adapted from de Barbaro et al., in press).

#### 2) How do tonic arousal levels change with increasing age?

During the first few years of life we transition from multiple shallow sleep-wake cycles through to more stable, once-per-day cycles (LeBourgeois et al., 2013). Tonic heart rate increases significantly from the 1st week to the 3rd month of life, and is stabilized at 6 months (Schwartz et al., 1989). It then decreases from typically 120 beats per minute (BPM) in 6-month-old infants to typically 70BPM in adults-although the exact development progression between infancy and childhood has not, to our knowledge, been tracked in detail.

Resting RSA is not typically observed in neonates, possibly due to inconsistencies in the periodicity of the respiration cycle at birth (Longin, Gerstner, Schaible, Lenz, & Koenig, 2006), but emerges over the course of the first year (Doussard-Roosevelt, Porges, Scanlon, Alemi, & Scanlon, 1997; Porges & Furman, 2011) and has reached adult levels by 5-7 years (Bornstein & Suess, 2000). Less change in RSA is observed among children and adolescents aged 8 and older (e.g. Quigley & Stifter, 2006; El-Sheikh, 2005).

Tonic pupil size is bigger in infants than in children (Wass, 2011), although again its exact developmental progression is unknown. Spontaneous fluctuations in pupil size increase with increasing age (Miller & Thompson, 1978). The number of spontaneous large-wave changes in EDA was found to increase during the first 10 weeks of life (Hernes et al., 2002) but the developmental progression of spontaneous EDA after this time has not, to our knowledge, been studied.

Research has identified spontaneous, periodic fluctuations in motor activity on a scale of minutes, that start in the fetal stage (Corner, 1977) and continue post-natally (Bacher & Robertson, 2001). Robertson found that these cyclic fluctuations decrease in a waking state during the first four months of post-natal life (Robertson, 1993), although they continue for longer in the sleeping state. Diurnal fluctuations start *in utero* and increase in amplitude during the first 3 months of life (Mirmiran & Lunshof, 1996), although the changes after that date have not to our knowledge been studied in detail. They have also been measured using actigraphs and heart rate recording (Mirmiran, Baldwin, & Ariagno, 2003) as well as cortisol (Seron-Ferre, Riffo, Valenzuela, & Germain, 2001).

#### 3) How do phasic changes in arousal relative to stimulus events change with increasing age?

Even young infants have been shown to demonstrate phasic ANS changes relative to experimenter-controlled stimulus events. For example, new-born infants show heart rate decelerations relative to low-intensity tones (Kearsley, 1973), although RSA changes are less obvious (Kearsley, 1973), and movement decelerations have been noted relative to new stimulus onsets (Byrne & Smith-Martel, 1987). EDA changes have also been demonstrated in 10-week-old infants in response to auditory stimuli (Hernes et al., 2002; see also Ham & Tronick, 2008). EDA responses to a heel prick increase in magnitude during the first month after birth (Storm, 2000)

Several researchers have shown that phasic ANS changes also increase in magnitude during the first year of life. Morrongiello and Clifton found that changes in HR relative to the presentation of a sound were more consistent in 5-month-old than in newborn infants (Morrongiello & Clifton, 1984). Byrne & Miller found that 6-month-olds showed a larger HR deceleratory response to the onset of a new stimulus than 3-month-olds, although no difference was found for body movement decelerations (Byrne & Miller, 1988). Hernes and colleagues found that the likelihood of infants showing a EDA response to an auditory stimulus increased from 8% to 50% between birth and ten weeks of age (Hernes et al., 2002).

Calkins and Keane found that between 2 and 4.5 years there was high stability in RSA suppression in response to challenge tasks at both ages. Modest cross-age stability in RSA suppression was also observed, along with a significant decrease in the magnitude of RSA suppression across age (Calkins & Keane, 2004).

Behaviorally, focused attention has also been reported to increase through the first year (Colombo & Cheatham, 2006). At birth, infants attend primarily to salient physical characteristics of their environment or attend with nonspecific orienting (Berg & Richards, 1997). Responsiveness to external stimuli develops rapidly through the first year, but voluntary, endogenous attention is generally thought to be almost completely absent until 12 months (Colombo & Cheatham, 2006). Richards examined the blink reflex, a response to high-intensity short-duration stimuli based on short-latency reflex pathways involving first-order neurons in the sensory pathways and brainstem cranial nerves (Balaban, Anthony, & Graham, 1989). Selective attention to one stimulus modality enhances the blink reflex to a stimulus of that modality and attenuates the blink reflex to stimuli in other modalities - so this attenuation of the blink reflex may also be used as an index of the amount of higher-order selective attention. Between 8 to 26 weeks, Richards found a clear increase in the enhancement of the blink reflex for the modality-match conditions and an increase in the attenuation of the blink reflex for the modality-mismatch conditions. This suggests an enhancement in selective attention across this age range (Richards, 2000).

In conclusion, research suggests that slow-moving changes in arousal covary with slow-moving changes in attention, as predicted by the AJM. No clear evidence is available on whether tonic arousal increases or decreases over development. However, phasic arousal changes relative to stimulus events and to attention both increase during the course of the first year.

### Part 2 - Infancy

In this section, we assess the predictions of the AJM for understanding infant development by asking the following three questions: first, behaviorally, is the best performance observed at intermediate levels of tonic (baseline) arousal? Second, are the largest phasic (reactive) changes observed at intermediate levels of tonic (baseline) arousal? Third, are larger phasic (reactive) changes associated with better performance? Separately, we then explain the fit of the model for understanding atypical development.

#### 1) Behaviorally, is the best performance observed at intermediate levels of tonic (baseline) arousal?

RSA is commonly used to index ‘tonic’ PNS control (lower arousal □ more PNS □ more RSA). Studies have suggested that 4-6-month infants with greater RSA (indexing greater PNS control) show better recognition memory (Frick & Richards, 2001; see also Linnemeyer & Porges, 1986). 3-6-month-old infants with high RSA look longer than low-RSA infants at a central stimulus in the presence of a distracting stimulus during sustained attention (Richards & Gibson, 1997; see also Richards, 1985a).

Two points should be made with respect to these findings. First, they have all documented linear relationships between arousal and attention (increased PNS □ decreased arousal □ better cognitive performance). This can be contrasted with the quadratic relationships predicted by the AJM, which predicts that hypo-, as well as hyper-arousal should both be associated with worse cognitive performance. The second point is that these papers only examined individual differences between individuals from the tonic (time-invariant) perspective. Given that both resting RSA and voluntary control of attention increase with increasing age, the findings reported could simply reflect that infants who were generally more developmentally advanced show both greater RSA and greater attention control, without there being a direct association between the two.

#### 2) Are the largest phasic (reactive) changes observed at intermediate levels of tonic(baseline) arousal?

The majority of studies in this area have measured RSA to index PNS control. Porges and colleagues found that newborn infants with high RSA responded to the onset of a stimulus tone with greater heart rate acceleration, and to the offset of the tone with greater heart rate deceleration (Porges, Arnold, & Forbes, 1973). A second study obtained similar findings when subjecting infants to changes in illumination (Porges, Stamps, & Walter, 1974). Other studies have also found that, in older infants, infants with higher RSA showed larger heart rate decelerations during sustained attention (Richards & Casey, 1991; Richards, 1987, Richards, 1985b).

Newborns with higher RSA also exhibit larger cortisol responses to a heel-stick procedure for drawing blood, suggesting greater stress reactivity (Gunnar, Porter, Wolf, Rigatuso, & Larson, 1995). In response to circumcision, newborn males with high RSA exhibited greater pain reactivity, as assessed by heart rate acceleration and fundamental cry frequencies (Porter, Porges, & Marshall, 1988).

Similar findings have been observed in research examining behavioral reactivity. For example, high RSA infants subjected to a pacifier withdrawal procedure cried more than their low RSA counterparts (Stifter & Fox, 1990). In a sample of premature infants DiPietro and Porges reported greater behavioral reactivity for high RSA neonates in response to a feeding procedure requiring a tube run through the nose or mouth (DiPietro, Porges, & Uhly, 1992).

As with the findings in the previous section, these findings point to linear relationships between arousal and phasic (reactive) changes, such that greater reactivity is observed at lower general levels of arousal (increased PNS □ decreased arousal ⍰greater reactivity).

#### 3) Are larger phasic (reactive) changes associated with better performance?

The AJM also predicts that greater phasic arousal changes could associate with better stimulus encoding. Across several studies Richards and colleagues have shown that greater (larger amplitude) HR decelerations, measured relative to both externally defined events (experimenter-defined stimulus presentations) and internally defined events (infants’ looks to and away from the target), index better stimulus encoding (reviewed Richards, 2010, 2011). For example, infants are better able to recognize material that was presented during phases of heart rate decelerations (Frick & Richards, 2001; Richards, 1997a). Infants are also less distractible during heart rate decelerations (Casey & Richards, 1988; Lansink & Richards, 1997).

Bornstein and Suess found that larger attention-related decreases in RSA (associated with a suppression of PNS activity in response to the presentation of a new stimulus) associated with more efficient habituation and with shorter looking (Bornstein & Suess, 2000). Similarly, DeGangi and colleagues found that greater RSA decreases during administration of the Bayley Scales associated with higher Mental Development Index scores in infants (G. A. DeGangi, DiPietro, Greenspan, & Porges, 1991). However, DiPietro and colleagues reported that infants who reacted to the stimulus with *increased* vagal tone (indicating phasic increases in PNS influence) were more often engaged in focused examination of an object (DiPietro et al., 1992). Richards and Casey discuss phasic *increases* in PNS influence during attention phases (which are the posited cause of the HR decelerations discussed above) (Richards & Casey, 1991). Thus, for RSA, convergent evidence suggests that increased phasic changes in RSA associate with better stimulus encoding. However, some of this research is inconsistent as to whether the relationship was one of an increase or a decrease in arousal relative to the stimulus event. (Recall that the AJM predicts that a larger phasic arousal change associates with better quality attention, but is agnostic as to whether the phasic change should be that of an increase, or a decrease in arousal.)

Wass and colleagues measured infants’ spontaneous visual attention (looks to and away from the screen) while presenting a stimulus battery consisting of mixed static and dynamic viewing materials to a cohort of typical 12-month-old infants (Wass, de Barbaro, Clackson, & Leong, under review). At the same time, they measured arousal via a composite of heart rate, galvanic skin response, head velocity and peripheral movement levels. They found that infants with generally more labile autonomic profiles showed more visual attention (longer look durations). Infants who showed more visual attention also showed greater phasic autonomic changes relative to attractive, attention-getting stimulus events. They also found that four sessions of attention training, which led to increased visual sustained attention, led to concomitant increases in arousal lability (Wass, de Barbaro, et al., under review).

Based on the same cohort as the Wass et al study, de Barbaro examined infants’ reactivity to a mild behavioral stressor. They found that infants who show high general levels of arousal reactivity show *lower* HR reactivity to a stressor (a video of another infant crying) (de Barbaro, Clackson, & Wass, 2016). Infants showing lower HR reactivity to a mild stressor showed a profile of reduced visual sustained attention but increased visual recognition memory.

Friedman and colleagues examined how body movement patterns during looking at 1-and 3 months relate to parent-reported attention problems at 8 years (Friedman, Watamura, & Robertson, 2005). They found that parent-reported attention problems at 8 years were associated with less suppression of body movement at the onset of looking, and greater rebound of body movement following its initial suppression at 3 months but not at 1 month (Friedman et al., 2005). This is consistent with a model that increased phasic ANS changes associate with better performance. Similarly, Robertson & Johnson examined the relationship between body movement changes during habituation and performance on that task (Robertson & Johnson, 2009). They distinguished between ‘suppressors’, for whom the typical decrease in body movement at the onset of looks persisted into the looks, and ‘rebounders’, for whom the initial decrease was more transient and movement quickly returns above baseline. Suppressors and rebounders did not differ on measures of looking during habituation, but when the stimulus changed rebounders looked more than suppressors.

In conclusion, research has shown that individuals who show increased phasic arousal changes to a sought-for stimulus show better voluntary control of attention and superior stimulus encoding. Recent findings have also shown that attention training, which increases infants’ visual attention (look durations to the screen) also leads to greater phasic changes in arousal profiles, suggesting a bidirectional effect.

### Atypical development

A number of studies have shown that early atypicalities in the auditory brainstem response can predict later atypicalities in aspects of social behavior and inhibition (Gardner et al., 2006; Geva, Schreiber, Segal-Caspi, & Markys-Shiffman, 2013; Geva, Sopher, et al., 2013). These atypicalities are, however, not ones of hypo-or hyper-arousal, but rather of inconsistent stimulus responses. Further, no research has investigated whether infants with atypical early brainstem responses show different patterns of phasic responses to experimenter-controlled stimuli during later development, as predicted by the AJM.

Along similar lines, Cohen and colleagues tested the auditory brainstem response as a prospective risk factor for subsequent development of Autism Spectrum Disorders (ASD) (Cohen et al., 2013; see also Miron et al., 2015). They found that children who had an early atypical brainstem response, together with a preference for high rates of visual stimulation at four months, were more likely subsequently to develop ASD (Cohen et al., 2013). A number of possible neural mechanisms have been discussed for the abnormal patterns of arousal-related change observed, involving reticular and thalamic areas (Cohen et al., 2013; Orekhova et al., 2012). Behaviorally, this may relate to findings from Wass and colleagues who found that 8-month-old infants who were later diagnosed with ASD showed more frequent eye movements, together with a reduced inability to modulate eye movement frequencies as a function of viewing time (Wass, Jones, et al., 2015). Behaviorally these behaviors are consistent with hyper-arousal, although arousal was not explicitly tested in this study.

In conclusion, although some research exists pointing to relationships between early arousal and subsequent atypicalities, no research has examined these questions from the perspective of the AJM.

### Part 3 - Childhood

In this section, we assess the predictions of the AJM model for understanding child development by asking the same three questions as in the previous section: first, behaviorally, is the best performance observed at intermediate levels of tonic (baseline) arousal? Second, are the largest phasic (reactive) changes observed at intermediate levels of tonic (baseline) arousal? Third, are larger phasic (reactive) changes associated with better performance? Separately, we then explain the fit of the model for understanding atypical development.

#### Typical development

##### 1. Behaviorally, is the best performance observed at intermediate levels of tonic (baseline) arousal?

Typical 3.5-year-old children with higher resting RSA (higher PNS ⍰ lower general arousal) show better performance on executive function tasks (Marcovitch et al., 2010). Similarly, in 6-13-year-old children, higher resting RSA associates with better performance on working memory and reaction time tasks (Staton, El-Sheikh, & Buckhalt, 2009). Higher resting RSA during infancy has also been shown to relate to better cognitive outcomes during childhood (Georgia A DeGangi, Porges, Sickel, & Greenspan, 1993). Suess and colleagues measured resting RSA in a cohort of typical 9-11-year-old children and recorded performance on the Continuous Performance Task (a measure of sustained attention) (Suess, Porges, & Plude, 1994). They found that higher RSA and lower rate associated with better performance on early (but not later) blocks of the task.

No research using methods other than RSA has, to our knowledge, addressed this question explicitly. However, some other indirect evidence is available. For example, some research has examined how circadian rhythms affect performance on cognitive tasks in children (van der Heijden, de Sonneville, & Althaus, 2010). The findings suggest that tasks with high voluntary (executive) demands show greater diurnal variability than tasks that are relatively more subserved by involuntary networks, such as pattern detection tasks (van der Heijden et al., 2010). However, arousal was not explicitly measured, and arousal changes in children as a function of circadian rhythms have not been tracked in detail (see Bonnemeier et al., 2003 for analogous work with adults).

Other indirect evidence comes from studies that used questionnaires to assess negative affect in children. Performance on voluntary attention tasks, such as a spatial conflict task, shows inverse correlations with negative affect in children of pre-school age (Gerardi-Caulton, 2000; Rothbart, Ellis, Rueda, & Posner, 2003 see also Aksan & Kochanska, 2004; Lawson & Ruff, 2004). Again, arousal was not directly measured in these studies, although other research has shown associations between arousal and questionnaire-based assessments of negative affect (Kolacz et al., 2016).

In conclusion, research has again identified linear relationships between arousal and performance, rather than the quadratic relationships predicted by the AJM. Better performance is shown in individuals at lower arousal (increased PNS). In addition, there is some evidence that certain tasks, putatively those with a larger ‘executive’ component, show larger variance with changing levels of arousal than others.

##### 2. Are the largest phasic (reactive) changes observed at intermediate levels of tonic(baseline) arousal?

A number of authors have reported linear relationships between baseline RSA and vagal withdrawal, such that higher RSA (lower arousal) is associated with increased withdrawal during a behavioral challenge in young children (Calkins, 1997; Marcovitch et al., 2010). However Suess and colleagues found no association between RSA and the amplitude of RSA decelerations during attention in children (Suess et al., 1994). To our knowledge no research using methods other than RSA has addressed this question.

##### 3. Are larger phasic (reactive) changes associated with better performance?

Again, the only research to have addressed this question in children has used RSA. Results are inconsistent in two aspects (see Graziano & Derefinko, 2013; Obradović & Finch, 2016 Porges, 2007). First, in the direction of effects: whether results show that phasic *increases* or *decreases* in RSA are considered to associate with ‘better’ performance. Second, in whether the results observed are linear (larger RSA change considered ‘better’) or quadratic (intermediate levels of RSA change associated with ‘better’ performance).

Greater RSA withdrawal in response to various laboratory challenges has been associated with increased sustained attention, engagement during challenge tasks, on-task behaviors in the classroom, and cognitive functioning, as well as more adaptive emotion regulation strategies (Blair & Peters, 2003; Calkins, Blandon, Williford, & Keane, 2007; Calkins & Keane, 2004; Doussard-Roosevelt, Montgomery, & Porges, 2003; Staton et al., 2009; Suess et al., 1994). Similarly, Becker and colleagues reported that better performance on an EF task was associated with greater RSA withdrawal among elementary school students (Becker et al., 2012). However, Utendale and colleagues found that greater EF skills were associated with *lower* RSA withdrawal among 5-and 6-year-olds, a relationship further moderated by children’s externalizing problems (Utendale et al., 2014). Staton and colleagues observed no relationship between RSA reactivity and cognitive performance in 6-13-year-old children (Staton et al., 2009). And Sulik and colleagues reported that while RSA withdrawal was linked to better EF performance, the associations between changes in RSA levels and EF skills varied across three different EF tasks (Sulik, Eisenberg, Spinrad, & Silva, 2015).

Studies described in the paragraph above have all documented linear relationships between RSA reactivity and performance. However, some other studies have observed quadratic relationships. For example, Marcovitch and colleagues observed a quadratic relationship in typical 3.5-year-old children, such that intermediate levels of RSA withdrawal during a number recall test, associated with better performance (Marcovitch et al., 2010). And ‘excessive’ RSA reactivity to challenges has been observed in psychiatric samples of internalizing and externalizing children, adolescents, and adults (Beauchaine, 2001; Beauchaine, Gatzke-Kopp, & Mead, 2007; Crowell et al., 2005; Mezzacappa et al., 1997).

Obradovic & Finch suggested than one reason for these inconsistent results may be that RSA studies tend to study change over relatively long time-scales (minutes) (Obradović & Finch, 2016). This may mean that a number of different subcomponent processes (initial withdrawal, maintenance of change, recovery) are all included in the measure of RSA withdrawal (Porges, 2007). By using piecewise growth curve modelling to examine change in RSA within the challenge period, they found that children with strong ‘cool’ EF skills showed curvilinear withdrawal during the challenge period, whereas those with lower levels showed curvilinear increases. But for ‘hot’ EF skills a different result was observed: those with higher EFs showed an inverted U-shaped trajectory (i.e. gradual, curvilinear RSA augmentation followed by mild RSA withdrawal), whereas those with lower hot EF skills displayed a U-shaped trajectory of RSA reactivity (i.e. gradual, curvilinear RSA withdrawal followed by mild RSA augmentation).

In conclusion, a number of studies have found that increased phasic (reactive) changes are associated with better performance. This is consistent with the AJM. However, other studies have suggested that ‘excessive’ phasic (reactive) changes are markers of psychopathology. This is inconsistent with the AJM, which suggests that increased phasic reactivity should be associated with better performance.

#### Atypical development

Atypical arousal has been noted in a number of aspects of child developmental psychopathology. In particular, lower RSA has been identified in a range of atypical populations, such as those with conduct disorder and depression (Pang & Beauchaine, 2013). Here, however, we concentrate on two well-researched clinical conditions. 1) Attention Deficit Hyperactivity Disorder, which has been suggested to involve tonic hyper-arousal, and 2) Autism Spectrum Disorders, which are thought to involve more complex patterns of atypical arousal.

##### 1) ADHD – tonic hyperarousal?

Children with Attention Deficit Hyperactivity Disorder (ADHD) move their heads more than typical children, and show more linear and less complex movement patterns (Teicher, Ito, Glod, & Barber, 1996; see also De Crescenzo et al., 2014). 24-hour analyses of heart rate also show that heart rate levels were overall higher in the ADHD group – with largest effects during afternoon and night (Imeraj et al., 2011). Findings for RSA are, however, inconsistent, with some authors reporting lower RSA in children with ADHD (Rash & Aguirre-Camacho, 2012) and others reporting no difference (Shibagaki & Furuya, 1997; Borger & van der Meere, 2000). Abnormal circadian patterns in children with ADHD have also been found using salivary cortisol (Imeraj et al., 2012). Although not completely consistent, these findings have led a variety of researchers to speculate that the pathophysiology of ADHD involves hyperactivation of the LC (Aston-Jones, Iba, Clayton, Rajkowski, & Cohen, 2007; Imeraj et al., 2012; Sonuga-Barke, Wiersema, van der Meere, & Roeyers, 2010) (although see also Clarke et al., 2013 for a contrasting approach).

Research into phasic arousal changes in children with ADHD has generally shown that children with ADHD show smaller phasic ANS changes relative to stimulus events. For example, Groen and colleagues examined HR decelerations during the performance of a selective attention task in which feedback was given. They observed enhanced HR decelerations in response to errors in typical children but not in those with ADHD. However, a Methylphenidate-treated ADHD group showed HR decelerations similar to those seen in typical controls (Groen, Mulder, Wijers, Minderaa, & Althaus, 2009). Boerger and van der Meere examined HR changes during Go/No-Go performance in children with ADHD and typical controls. They found no evidence of decreased HR decelerations relative to the onset of the No-Go signal, but they did find that heart rate decelerations before the onset of Go signals, which are believed to reflect motor preparation, were less pronounced in the ADHD children (Borger & van der Meere, 2000; see also Shibagaki & Furuya, 1997). Using a similar model, O’Connell and colleagues examined EDA responses to errors in children with ADHD. They found reduced EDA responses in the ADHD group, and that sustained attention errors were predicted by EDA amplitudes (O’Connell, Bellgrove, Dockree, & Robertson, 2004).

##### 2) ASD – atypical arousal?

Autism Spectrum Disorders (ASD) may show a more complex pattern. There are some reports of hyper-tonicity similar to that suggested for ADHD. For example, Bal and colleagues found a faster tonic heart rate in children with ASD (Bal et al., 2010; see also Heilman, Harden, Zageris, Berry-Kravis, & Porges, 2011). They also reported lower amplitude RSA (suggesting decreased PNS contribution) (although Watson, Roberts, Baranek, Mandulak, & Dalton, 2012 failed to replicate this). Anderson and Colombo noted increased tonic pupil size in a small sample of 23-70-month-old children with ASD (Anderson & Colombo, 2009). This can be compared to studies identifying increased brain metabolic activity at rest in individuals with ASD, suggesting a failure to deactivate at rest (Kennedy, Redcay, & Courchesne, 2006).

Schoen and colleagues examined phasic EDA changes in children with high-functioning ASD in response to presentation of mixed sensory stimuli. They identified two subgroups-a hyper-aroused group that showed increased attention-related changes and faster latencies, and a hypo-aroused subgroup that showed lower changes and slower latencies (Schoen et al., 2008, see also Hirstein, Iversen, & Ramachandran, 2001). Ben-Sasson and colleagues noted strikingly similar results in a behavioral analysis looking at sensory sensitivities (Ben-Sasson et al., 2008; see also Leekam, Prior, & Uljarevic, 2011). They reported co-occurring auditory hyper-and hypo-sensitivity symptoms in children with ASD, and suggested that both of these problems may be explained by a common mechanism, namely a dysfunctional arousal system that compromises the ability to regulate an optimal response (Ben-Sasson et al., 2008). Schaff and colleagues found that, whereas typical children showed a decrease in RSA in response to challenging stimuli, children with ASD did not (Schaff et al., 2013).

In conclusion, results from children with ADHD are largely consistent with the predictions of the AJM. Children with ADHD have been suggested to show hyper-tonic activation, together with reduced phasic responsiveness and poorer top-down control of attention. Results from children with ASD are thought to be more complex, with the possibility of hyper-and hypo-aroused subgroups.

## Discussion

### How well does the evidence fit the Aston-Jones model?

We have reviewed the relationship between: 1) an individual’s level of tonic (baseline) arousal; 2) the degree of phasic (reactive) change they show relative both to sought-for stimulus events and to mild behavioral challenges; and 3) levels of cognitive performance.

The Aston-Jones model (AJM), following on from Yerkes and Dodson, predicts that a quadratic relationship should be observed between arousal and performance, such that best performance is seen at intermediate levels of arousal. It also states that the largest degree of phasic (reactive) change should be observed at intermediate levels of tonic (baseline) arousal.

The majority of the evidence that we have reviewed shows consistent relationships between tonic arousal, phasic arousal changes and performance – but it points to linear relationships between the three variables. Behaviorally, best performance tends to be observed in individuals at *lower* levels of tonic arousal; larger phasic changes are observed in individuals at *lower* tonic arousal; and better performance is observed in individuals who show *larger* phasic arousal changes. This pattern is equivalent to the right-hand side of the inverted U-relationship shown in Figure 1. With the exception of one study (from Schoen and colleagues, looking at ASD (Schoen et al., 2008)), no research has documented, or examined, the hypo-aroused phenotype, predicted by the AJM, and Yerkes-Dodson.

There are a number of possible explanations for this. The first is that the majority of the research, particularly with children, has used a single measure, RSA, to quantify arousal, and it is unclear whether RSA is sensitive to hypo-as well as hyper-arousal. It may be that other research using, for example, movement patterns, might be more sensitive to both extremes of arousal. The second is that infants and children may have a tendency, relative to adults, towards hyper-tonic arousal. (A finding not directly empirically investigated (see Part 1), but to which many parents of young children would testify.) What is in effect a normal curve thus becomes positively skewed, making the positive linear trend stronger than the quadratic effect. A third possibility is that the specific demands of testing (attending a lab, wearing equipment) or excluding trials in which no response was given mean that hyper-arousal is over-represented relative to hypo-arousal in experimental samples, and that hypo-arousal might be observed more in naturalistic contexts (such as the classroom) than it is in the lab. A fourth possibility is that the quadratic effect exists but studies are underpowered to recognize it.

One further challenge for the AJM is that the three elements it connects – tonic (baseline) arousal, phasic (reactive) arousal changes and performance – are treated as showing unitary co-variance. However it may be that the influence of arousal on performance is stronger in some individuals than others (Breeden, Siegle, Norr, Gordon, & Vaidya, 2016). Metin and colleagues, for example, examined how changing event rate (the rate of stimulus presentation) affects performance both in typical children and in children with ADHD (Metin et al., 2013). (Event rate was used as a proxy for arousal, since fast-paced stimuli were considered to be more arousing.) Results showed that altering the event rate (‘increasing arousal’) had a larger effect on performance in children with ADHD than in typical children, which they interpreted as suggesting that performance in children with ADHD may be disproportionately affected by changes (either increases or decreases) in arousal. Similarly we might predict that, due to asymmetric patterns of brain maturation (with subcortical attention networks developing before cortical attention networks) (Johnson, Posner, & Rothbart, 1991; Colombo, 2001), arousal-attention interactions might be stronger in infants than in older children. If proven, such a finding would be hard to incorporate into the AJM.

### Implications for understanding Differential Susceptibility Theory

In the Introduction, we asked: do the same individuals who show a large arousal change to a negative stimulus (mild experimental stressors such as arm restraint) also show a large arousal change to a positive stimulus (such as a new item to be memorized in a working memory task, or a novel image in a habituation task)?

Some of the research we have presented is consistent with this model. For example, newborn infants with high levels of RSA show increased reactivity to negative events, such as a heel stick procedure for drawing blood (Gunnar et al., 1995; see also Stifter & Fox, 1990; Porter et al., 1988). And they also show increased reactivity to a sought-for stimulus event, or during phases of sustained attention (Richards & Casey, 1991; Richards, 1985b). Thus, RSA may reflect the capacity to engage in both a positive, and a negative, manner (Beauchaine, 2001).

These findings are also consistent with other research examining long-term associations of RSA. For example, elevated RSA reactivity during a parent–child interaction was associated with less optimal performance on executive function tasks in 3- to 5-year-olds exposed to maltreatment, and with more optimal performance among nonmaltreated children (Skowron et al., 2014). And in the context of poor family circumstance, 5-6-year-old children with high RSA reactivity showed the worst outcomes, whereas children with low RSA reactivity showed the best outcomes (Obradovic et al., 2010; see also Peltola et al., 2016).

This suggests that individuals with high RSA would show increased phasic (reactive) autonomic changes in response both to positive and negative events. These increased phasic (reactive) changes would confer performance advantages in ‘positive’ situations, where the stimulus is interesting or sought-for. However, they would also confer disadvantages in situations in which the stimulus is unexpected, intense or aversive. This is consistent with our suggestion that the AJM may offer a mechanism for understanding the short-term interactions between neurobiological sensitivity and cognitive performance, which is consistent with DST.

However, not all of the research we have reviewed is consistent with this framework. The primary inconsistency is research pointing to *excessive* phasic (reactive) RSA changes as a risk marker for psychopathology (Marcovitch et al., 2010; Pang & Beauchaine, 2013). The idea that excessive reactivity may be a risk factor for psychopathology relates to neurochemical evidence suggesting that chronic stress enhances the excitability of the HPA and sypatho-adrenomedullary systems (Ulrich-Lai & Herman, 2009). But it is inconsistent with the AJM, which points to larger phasic change as a marker of better performance, and suggests that greatest reactivity should be observed in individuals at intermediate levels of tonic (baseline) arousal.

One likely reason for this is that research findings are being integrated across studies that have explored phasic (reactive) change across multiple time-scales. Aston-Jones recorded phasic changes to stimuli on a time-scale of milliseconds, whereas most studies examine RSA reactivity over a time-scale of minutes. As a number of authors have noted, recording RSA reactivity over long time-periods means that distinct physiological processes of reactivity and recovery contribute to the reactivity measure (Obradović & Finch, 2016; Schmitz, Krämer, Tuschen-Caffier, Heinrichs, & Blechert, 2011). Consequently, researchers have called for more dynamic analytic approaches to investigating individual differences in physiological response trajectories (Burt & Obradovic, 2013; Wass, Clackson, & Leong, under review; Obradović & Finch, 2016). One possibility might be that atypical or high-risk populations, who are known to show lower tonic (baseline) RSA, might show small initial reactivity combined with a failure of subsequent regulatory/compensatory processes. Examining change on a more fine-grained time-scale will allow us to address these questions.

One further challenge is that the majority of work has failed to distinguish between different types of behavioral challenge – despite that individuals show differing levels of sensitivity to different types of challenge (Obradović, Bush, & Boyce, 2011; Quigley & Stifter, 2006; Sulik et al., 2015). More research is needed that explicitly addresses this issue – and, in particular, directly addresses the question with which we started, which is that of whether individuals who show large arousal changes to a ‘negative’ stimulus (mild experimental stressors such as arm restraint) also show large arousal changes to a ‘positive’ stimulus (such as a new item to be memorized in a working memory task, or a novel image in a habituation task). Relatively little of the research that we have covered in this review has directly addressed this question, within a single cohort.

### Directions for future work

We have also noted a variety of gaps in the literature, and directions for future work. Specifically:

i) one area that has received relatively little attention is the hypo-aroused subgroups predicted by the Yerkes-Dodson and Aston-Jones frameworks. The failure to identify these subgroups may reflect that the majority of research in this area has used RSA, which may not be sensitive to hypo-arousal. Future research using other methods such as actigraphy should explore this in more detail.
i) future work should explicitly examine change on multiple different time-scales. Of particular interest is to compare studies that examined reactivity at the millisecond-level scale with those that examine reactivity over longer time-scales. Conducting this research is essential in order to allow us to distinguish reactivity from other processes such as maintenance of change and recovery.
i) Previous research has suggested that, whereas voluntary, ‘top-down’ attention control is highest during mid-level arousal, some aspects of memory encoding should be highest during hyper-arousal (Arnsten, 2009; de Barbaro et al., 2016; Wass, de Barbaro, et al., under review). Future research should explore in more detail the strengths, as well as the weaknesses, of the hyper-aroused behavioral phenotype.
i) finally, the implications for intervention of this work remain to be investigated. The AJM predicts that different tactics should be employed contingent on the state of the child: in a hyper-aroused child, decreasing arousal should associate with increasing performance, whereas in a hypo-aroused child the opposite relationship should be observed. However, these types of predictions arising from the model remain, to our knowledge, almost completely unexplored.

## Acknowledgements.

This article was supported by a British Academy Postdoctoral Fellowship, an ESRC Future Research Leaders Fellowship (ES/N017560/1) and by intramural funding at the Medical Research Council Cognition and Brain Sciences Unit at Cambridge. Thanks to Kaya de Barbaro, Roeljan Wiersema and Edmund Sonuga-Barke for useful discussions.

